# Myocardial Endoglin Regulates Cardiomyocyte Proliferation and Cardiac Regeneration

**DOI:** 10.1101/2024.09.27.615380

**Authors:** Daniel W. Sorensen, Philip M. Tan, Bayardo I. Garay, Jacob Solinsky, Doğacan Yücel, Michael J. Zhang, Conor Galvin, Henry Elsenpeter, Jennifer Mikkila, Kendra Jerdee, Timothy D. O’Connell, Jeffery D. Molkentin, Rita C.R. Perlingeiro, Yasuhiko Kawakami, Jeffrey J. Saucerman, Jop H. van Berlo

## Abstract

The mammalian heart loses almost all its regenerative potential in the first week of life due to the cessation of the ability of cardiomyocytes to proliferate. In recent years, a number of regulators of cardiomyocyte proliferation have been identified. Despite this, a clear understanding of the regulatory pathways that control cardiomyocyte proliferation and cardiac regeneration is lacking, and there are likely additional regulators to be discovered. Here, we performed a genome-wide screen on fetal murine cardiomyocytes to identify potential novel regulators of cardiomyocyte proliferation. Endoglin was identified as an inhibitor of cardiomyocyte proliferation in vitro. Endoglin knock-down resulted in enhanced DNA synthesis, cardiomyocyte mitosis and cytokinesis in mouse, rat and human cardiomyocytes. Using gene-targeted mice, we confirmed myocardial Endoglin to be important in cardiomyocyte proliferation and cardiac regeneration using gene-targeted mice. Mechanistically, we show that Smad signaling is required for the endoglin-mediated anti-proliferative effects. Our results identify the TGF-β coreceptor Endoglin as a regulator of cardiac regeneration and cardiomyocyte proliferation.

**Summary:** High-content function screening is used to identify a novel inhibitor of cardiomyocyte proliferation which can promote mammalian cardiac regeneration.

## Introduction

One of the most important reasons for the devastating outcomes of heart disease is the lack of regenerative potential of the adult heart (Van Berlo et al., 2014; Virani et al., 2020). Both acute and chronic heart diseases are accompanied by a loss of myocardium and a related loss of contractile performance (Del Re et al., 2019; van Empel et al., 2005). This loss of myocardium is not compensated for by an increase in new cardiomyocyte (CM) generation. It is now well-established that the adult heart does not harbor a dedicated progenitor cell population and has an extremely low level of adult CM turnover (Bergmann et al., 2009; Kimura et al., 2015; Senyo et al., 2013; Van Berlo et al., 2014). The lack of repair of the adult heart in response to an injury is in stark contrast to the cardiac regeneration that is observed throughout animal diversity, such as in zebrafish, urodele salamanders and even in neonatal mice (Oberpriller and Oberpriller, 1974; Porrello et al., 2011; Poss et al., 2002; Raya et al., 2003). The common feature in repair of the cardiac injury in these animals is a reactivation of the proliferative potential of CMs (Kikuchi et al., 2010; Mohamed et al., 2018).

Induction of CM proliferation presents several unique challenges. CMs are terminally differentiated, with many cells having tetraploid DNA content. This is likely an important reason for the relative refractoriness of adult CMs to become proliferative (Patterson et al., 2017). Nevertheless, manipulating core cell cycle regulators has shown promise to induce adult CMs into a proliferative state (Cheng et al., 2007; Mohamed et al., 2018; Pasumarthi et al., 2005; Porrello et al., 2011). Some of the most potent and convincing evidence for reactivation of cell cycle and completion of cell division in adult CMs came from induced expression of 4 separate cell cycle regulators, or alternatively, 2 cell cycle regulators in conjunction with Wee1 and TGF-β inhibition (Mohamed et al., 2018). Additional regulators of CM proliferation have also been identified such as Nrg (Bersell et al., 2009; Eulalio et al., 2012). Another strong and convincing regulator of CM proliferation comes from interference with the Hippo signaling pathway. Genetic deletion of the Hippo regulator Salvador from CMs in mice resulted in hyperplastic hearts with excessive heart size for the weight of the mouse (Heallen et al., 2011).

Despite insights into specific regulators of CM proliferation, little is known about the overall regulatory signaling that governs cell cycle exit in cardiomyocytes soon after birth. The investigation into CM proliferation has been largely limited to known cell cycle regulators which have well established roles in other cell types, such as expression of cyclin-dependent kinases, under control of Retinoblastoma proteins (Di Stefano et al., 2011; Porrello et al., 2011). Here, we performed an unbiased genome-wide screen on primary fetal murine CMs to identify additional genes that regulate cardiomyocyte cell cycle exit. Using high-content imaging after gene knockdown, we screened approximately two-thirds of the protein-coding genes in the mouse genome for their effect on cardiomyocyte proliferation. New potential regulators of cardiomyocyte proliferation were identified, although some of these were insufficient to inhibit proliferation in-vivo. We identified Endoglin (*Eng*), which encodes a TGF-β coreceptor, as an inhibitor of cardiomyocyte proliferation. We show that depletion of *Eng* from cardiomyocytes but not from cardiac fibroblasts, is sufficient to promote functional regeneration of the myocardium after injury, via cardiomyocyte proliferation. Mechanistically, we show that Eng inhibits cardiomyocyte proliferation in a Smad2-dependent manner. These results identify a novel mechanism regulating cardiomyocyte proliferation and myocardial regeneration in mammals.

## Results

### Screen to Identify Novel Inhibitors of Cardiomyocyte Proliferation

To identify novel inhibitors of CM proliferation we performed a genome-wide screen using lentiviral shRNA particles to knock-down individual genes, in fetal murine CMs, isolated from E17.5 mice, a stage where CMs are still proliferative in vivo (Soonpaa et al., 1996). We used Myc overexpression as a positive control, since Myc is known to induce CM cell cycle activation (Bywater et al., 2020; Muñoz-Martín et al., 2019). Lentiviral scrambled shRNA expression was used as a negative control. To identify CM cell cycle activation, we added the thymidine analog 5-Ethynyl-2’-deoxyUridine (EdU) for the last 24h of the culture period, followed by staining the fixed cells for incorporated EdU and for troponin T, to identify CMs. We utilized a previously published lentiviral shRNA library, where individual shRNAs are used with an average of 4-5 shRNAs per gene for a total of ∼15,500 genes (Moffat et al., 2006). We imaged 18 sites per well, accounting for >6 million images that were analyzed (Figure 1A). We optimized our previously developed pipeline for high-content screening of CM cultures to quantify proliferative vs non-proliferative CMs (Bass et al., 2012; Sutcliffe et al., 2018)(Figure 1B and Supplemental Figures 1A and 1B). All positive wells that were identified by the automated algorithm were manually screened and verified. After manual verification we identified 251 genes as potential negative regulators of CM proliferation (Figure 1C). To verify the validity of this primary screen, which was performed with an n=1, we generated new lentiviral particles for all shRNAs that induced CM proliferation and rescreened these to verify that they also stimulated DNA synthesis in CMs. Of the initial 251 putative regulators of CM proliferation, we verified 146 to enhance EdU incorporation in CMs (Supplemental Table 1). These included well-known regulators of cell cycle activity, such as Cdkn1a and Rbl1(Di Stefano et al., 2011). In addition, we identified Mapre3 and Rab8a as potential regulators of CM proliferation based on enhanced EdU incorporation (Figure 1D, Supplemental Table 1) These genes had previously been linked to cell cycle progression (Ferreira et al., 2013; Liu et al., 2022). To further explore the possibility that identified hits from the primary screen might indeed be regulators of CM proliferation, we stained cultured CMs for phosphorylated Histone H3 (pHH3), a marker of progression to the G2/M phase of cell cycle. We noted enhanced pHH3 staining in some, but not all conditions, with shRNA targeting Cdkn1a and Rab8a clearly showing pHH3 staining, while Rbl1 and Mapre3 had barely any CMs showing pHH3 staining (Figure 1E). When we plotted the level of pHH3 staining of individual wells for the top 40 genes, many of the EDU+ cells failed to show a corresponding increase in pHH3 staining (Figure 1F). Regardless, there was high correlation between EdU incorporation rates and pHH3 staining levels, suggesting that stimulation of EdU incorporation typically does result in enhanced pHH3 staining levels in these primary fetal CM cultures (Figure 1G and Supplemental Table 1). Interestingly, some of the most potent inhibitors of CM proliferation in-vitro did not appear to regulate CM proliferation in-vivo. Knocking down Mapre3 in-vitro produced a robust increase in the frequency of in S-phase progressing CMs (Figure 1D, Supplemental Table 1). However, the genetic loss of myocardial Mapre3 in-vivo, did not significantly alter the rate of CM proliferation in mice following an apical resection injury (Supplemental Figure 2A-B). Additionally, Cdkn1a knockout (KO) mice did not show a significant change in the frequency CMs that were positive for pHH3 after injury (Supplemental Figure 2C-D). In the Cdkn1a knockout mice, an upregulation of Cdkn1b could potentially explain the lack of enhanced proliferation since combined knockdown of Cdkn1a and Cdkn1b in vivo has previously been shown to induce CM hyperplasia (Di Stefano et al., 2011). It is possible that some of the identified inhibitors are only sufficient to inhibit CM proliferation in vitro and their depletion needs to be paired with others to promote proliferation in vivo.

**Figure 1.**
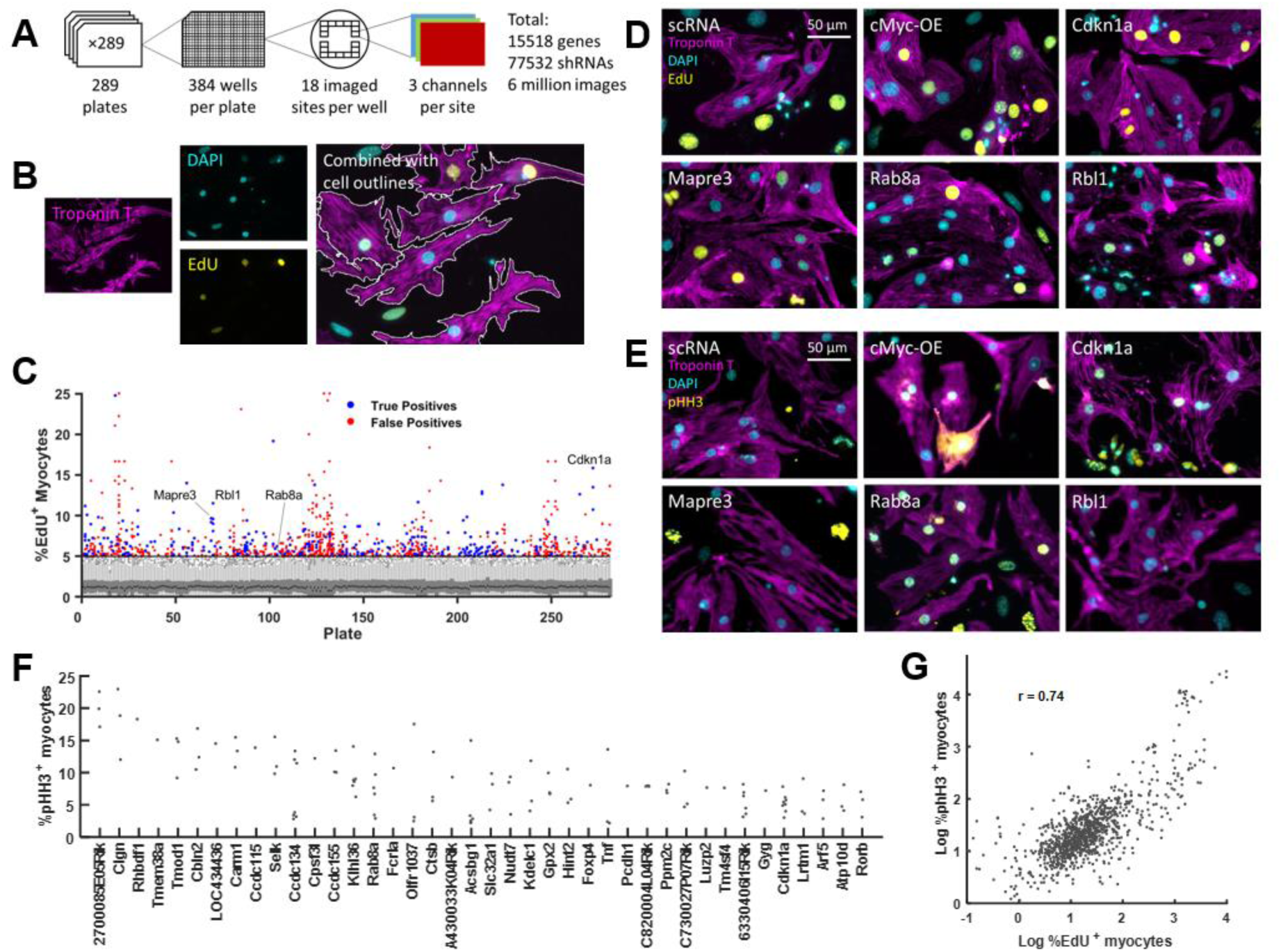
Genome-wide Screen to Identify Inhibitors of CM Proliferation. (A) Schematic overview of genome-wide shRNA screen. (B) Example of automated detection of proliferative (EdU+) CMs (TnT+). (C) Manhattan plot showing results of primary screen showing positives in blue (True positives) or red (False positives). Select genes are identified by name. (D) Representative images of EdU (yellow) incorporation for selected genes from confirmation screen. (E) Representative images of pHH3 staining for selected genes from confirmation screen. (F) Scatter plot showing level of pHH3 positivity for top 40 genes. (G) Correlation plot showing correlation between EdU positivity and pHH3 staining.

### Endoglin Inhibits Mammalian Cardiomyocyte Proliferation

Our interest was piqued by one of the CM regulators identified from the screen, Endoglin, which was one of the few genes had previously been linked to myocardial development. Mice with a germline deletion of the TGF-β coreceptor Eng have vascular defects which causes their premature demise at mid-gestation, but these mice also displayed dilated hearts (Arthur et al., 2000). In order to see if Eng was physiologically meaningful later in development we compared cardiomyocyte mRNA expression early in postnatal development at day 1, when mice are capable of robust CM proliferation and later when that ability is lost at adulthood. *Eng* mRNA levels in CMs increased from a neonatal age to adulthood correlating with the loss of regenerative capacity, suggesting that Eng might be contributing to the diminishment of cardiac regenerative potential (Figure 2E). This relationship appears to be conserved in humans as well. Eng mRNA in human ventricles was significantly enriched in adults compared to fetuses (9-28 weeks old) (Supplemental Figure 2E). Although Eng was not the most potent inhibitor of CM proliferation we identified, lentiviral knockdown resulted in a robust and significant increase in the fraction of CMs progressing to S-phase and G2/M-phase of the cell cycles (Figure 2B-E). To ensure that the observed activation of CM cell cycle was conserved among species and later in development, we treated Neonatal Rat Ventricular Cardiomyocytes (NRVM) with scrambled siRNAs and compared this with *Eng* targeting siRNAs. We first verified knockdown of Eng from NRVMs (Supplemental Figure 2F). We then added EdU to NRVMs for the last 24h of the culture, followed by fixation and staining for incorporated EdU and Troponin T to identify proliferative CMs. We observed a significant enhancement of the fraction of NRVMs that were progressing through S phase of the cell cycle (Figure 2F-G). We next verified that NRVMs with Eng knockdown would progress to G2/M phase of the cell cycle by staining for pHH3, and indeed observed a larger fraction of NRVMs that stained positive for pHH3 after Eng knockdown (Figure 2H-I). To further assess if NRVMs would undergo cytokinesis, we stained for the cleavage furrow and cytokinesis marker Mklp1(Powers et al., 1998), which was not observed in scrambled siRNA treated NRVMs, but observed in *Eng* siRNA treated NRVMs (Figure 2J and Supplemental Figure 2G). These results indicate that *Eng* knock-down is sufficient to stimulate mouse and rat CM proliferation in vitro.

**Figure 2.**
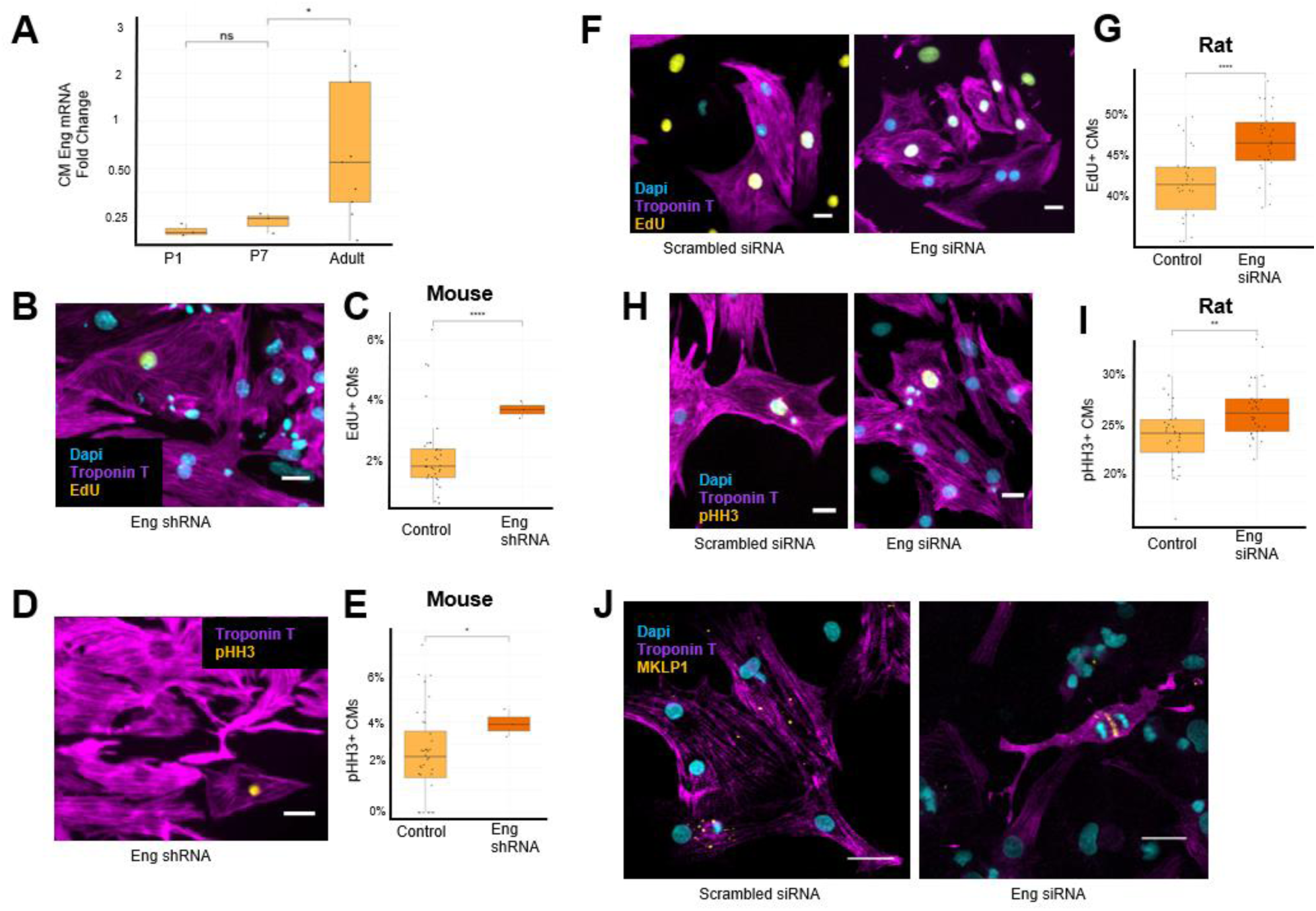
Eng Inhibits Mammalian CM-Proliferation in vitro. (A) Relative Eng mRNA expression in CMs at indicated postnatal developmental time points (B) Representative image of Eng shRNA treatment showing EdU incorporation in fetal murine CMs (C) Quantification of EdU positive CMs after treatment with scrambled or Eng shRNA laden lentivirus (D) Representative image of Eng shRNA treatment showing pHH3 and Troponin T staining (E) Quantification of pHH3 positive fetal murine CMs after treatment with scrambled or Eng shRNA (F) Representative images of scrambled or Eng siRNA treatment showing EdU incorporation in NRVMs (G) Quantification of EdU positive NRVMs after treatment with scrambled or Eng siRNA (H) Representative image of scrambled or Eng siRNA treatment showing pHH3 staining in NRVMs (I) Quantification of pHH3 positive CMs after treatment with scrambled or Eng shRNA (J) Mklp1 and Troponin T staining of NRVMs after treatment with scrambled or Eng siRNA.

### Myocardial Endoglin Inhibits Cardiomyocyte Proliferation After Cardiac Injury

We next determined whether Eng was able to regulate CM proliferation in vivo in the context of neonatal myocardial injury. To accomplish this, we cross-bred a loxP targeted Endoglin (Eng^fl/fl^) mouse strain with a tamoxifen-inducible cardiomyocyte-Cre strain (Myh6^mer-cre-mer/+^) to generate mice that would allow CM-specific deletion of Eng in an inducible manner (Allinson et al., 2007; Sohal et al., 2001). We used Eng^fl/fl^ X Myh6^mer-cre-mer/+^ (CM Eng KO) as experimental mice and Myh6^mer-cre-mer/+^ (Control) as control mice. Both experimental and control mice were administered tamoxifen one day after birth, followed by apical resection at seven days after birth (Mahmoud et al., 2014; Sadek et al., 2014). To quantify the number of CMs that entered S-phase we administered EdU for 5 consecutive days, starting at 6 days post-apical resection (Figure 3A). Ventricles were harvested at 21 days after birth, and we quantified EdU-positive, PCM1-positive ventricular CM nuclei. CM Eng KO mice showed enhanced EdU incorporation compared with controls (Figure 3B-D). Moreover, CM Eng KO mice harvested 7 days after injury showed increased frequencies of pHH3-positive CMs indicative of CMs progressing to mitosis (Figure 3C-E). These findings were further supported by Aurora Kinase B staining in CM Eng KO mice after apical resection, indicative of active CM cytokinesis (Figure 3F). To determine if the enhanced proliferative response to injury in CM Eng KO mice would result in improved heart regeneration, we assessed cardiac function after injury by echocardiography. Indeed, CM Eng KO mice displayed improved cardiac function compared with control mice 38 days after injury, as demonstrated by increased ejection fraction and fractional shortening (Figure 3G-H and Supplemental Figure 3A). No significant functional difference was observed between genotypes in uninjured animals (Supplemental Figures 3B,C). To further assess if CM-specific deletion induced cardiac regeneration, we measured overall heart size after injury, which showed no difference between CM Eng KO and controls (Supplemental Figure 3D). Histological examination showed that CM Eng KO hearts displayed a reduced area of fibrosis at the injury site, suggesting a replacement of the scar tissue by regenerated myocardium (Figure 3I-J). These results demonstrate that CM-specific deletion of Eng is sufficient to promote CM proliferation and functional myocardial regeneration after neonatal myocardial injury.

**Figure 3.**
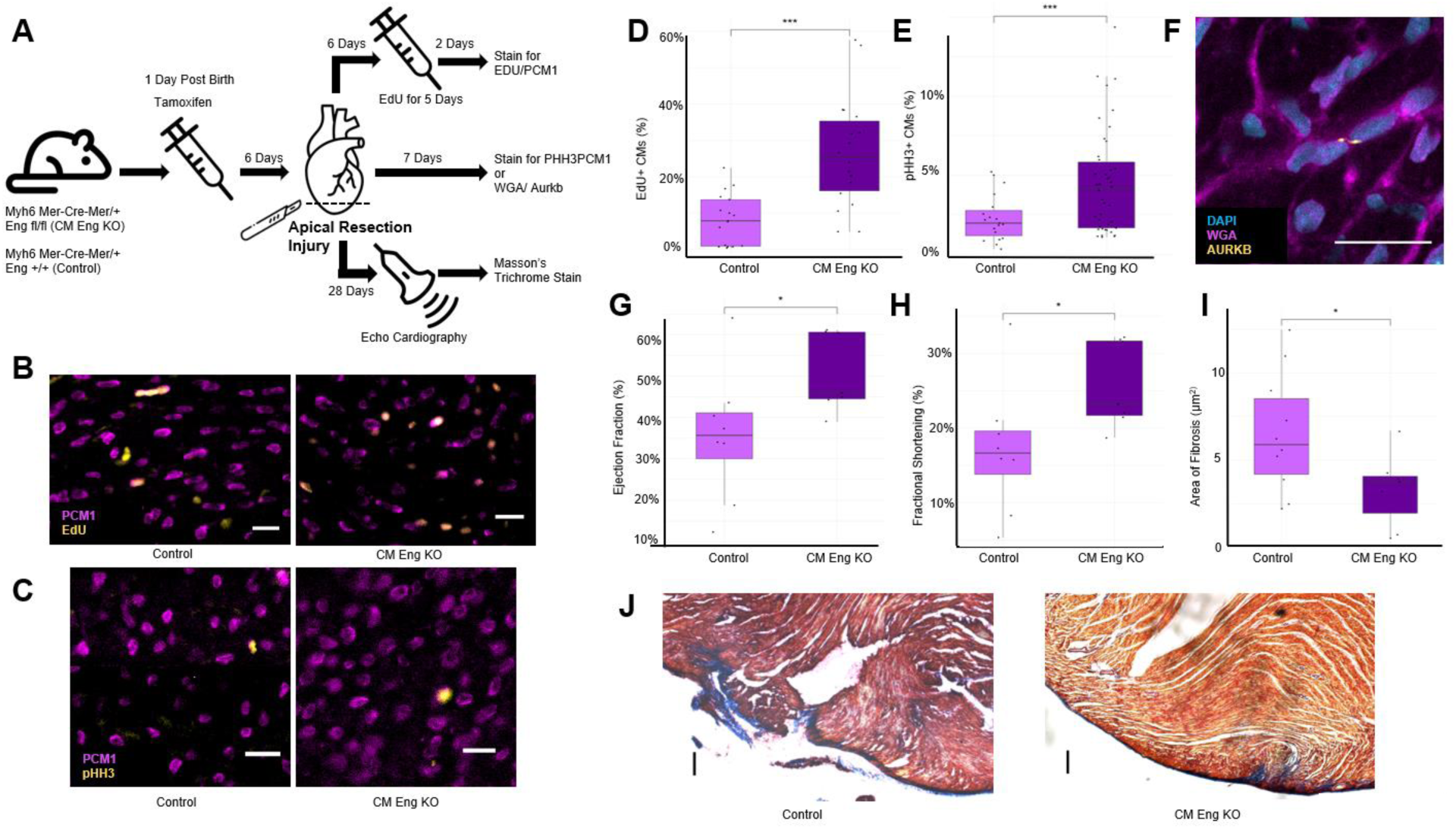
Myocardial Eng Knockout Results in Cardiomyocyte Proliferation and Cardiac Regeneration. (A) Schematic of control and experimental animals and experimental timeline of the conditional myocardial Eng-KO line (B) Representative images of EdU and Pcm1 staining after apical resection to detect DNA synthesis in CMs (C) Representative images of pHH3 and PCM1 staining after apical resection (D) Quantification of control and CM Eng-KO EdU+ CMs 13 days after Injury, N=17/17 (E) Quantification of pHH3+ CMs in control and CM Eng-KO mice 7 days after injury, N=18/41 (F) Example WGA, DAPI and Aurora B Kinase staining labeling of a cytokinetic event in a CM Eng-KO ventricle (G) Ejection Fraction and (H) Fractional Shortening of control and CM Eng-KO mice 38 days after Injury N=8/7 (I) Representative Masson’s Trichrome Staining from control and CM Eng KO mice 39 days after injury. (J) Quantification of fibrosis of control and CM Eng KO mice 39 days after injury, N=10/7.

### Fibroblast Endoglin is Not Responsible For Eng-Mediated Inhibition of Cardiomyocyte Proliferation

Endoglin is expressed in the heart, but CMs are only a minor source of Eng expression (Figure 4A). It has been shown that cardiac fibroblasts are a major cell type to produce Eng in the ventricle (Kapur et al., 2012). Eng is initially produced as a membrane-bound TGF-β coreceptor. However, Eng can also be cleaved by Matrix Metalloproteinase 14 to produce soluble Eng (sEng) (Tu’uhevaha et al., 2012). These different forms of Eng have been shown to have distinct mechanism of signaling and effects on the cell cycle (Núñez-Gómez et al., 2017). The transmembrane form (mEng) acts as a TGF-β coreceptor and can be phosphorylated by activated receptor complexes. sEng has been proposed to competitively sequester TGF-β and BMP ligands from their receptors, but can also form functional sEng/Bmp ligand complexes (Lawera et al., 2019). Since fibroblasts are a major source of Eng in the heart, fibroblast Eng could be solubilized to potentially mediate the proliferative effect on CMs. We next determined the effect of fibroblast-specific Eng expression on CM proliferation and cardiac regeneration. To this end, we cross-bred the loxP targeted Eng mouse line with a tamoxifen-inducible fibroblast-Cre expressing mouse (Tcf21^mer-cre-mer/+^, Fibro Eng KO) (Figure 4B) (Acharya et al., 2011). This strategy preserves CM-specific expression of Eng and allows for assessment of the function of fibroblast-derived Eng, including sEng. We subjected Fibro Eng KO and control animals to the same injury model and timeline as described in Figure 3A. Similar to CM Eng KO, we noted an increase in EdU+ CMs in Fibro Eng KO animals, compared with controls, although the magnitude of enhanced S-phase progression in CMs was not as high as in CM Eng KO mice (Figure 3D, Figure 4C-D). However, Fibro Eng KO animals displayed no significant change in pHH3 positivity (Figure 4E-F). These results suggest that depletion of fibroblast-derived Eng does not result in increased CM mitosis. To determine if this incomplete stimulation of CM cell cycle activation was sufficient to enhance cardiac regeneration, we again measured cardiac function after apical resection. Consistent with lack of cell cycle progression of CMs to the G2/M phase of cell cycle, we observed no improvement in cardiac function in Fibro Eng KO animals compared with controls (Figure 4G-H, Supplemental 4A-B). No change in heart weight vs body weight ratios was observed (Supplemental Figure 4C). Interestingly, we noted a substantial reduction in scar tissue in Fibro Eng KO mice compared with controls, suggesting that lack of scar formation is insufficient to enhance CM proliferation and cardiac regeneration (Figure 4I-J). To further validate our in vivo findings, that fibroblast-derived sENG does not modulate CM proliferation, we utilized TRC105, an antibody that can specifically block sEng’s ability to bind to key ligands such as Bmp9 (Liu et al., 2021; Nolan-Stevaux et al., 2012). Application of the antibody alone did not modulate CM proliferation (Figure 4K). Furthermore, application of both Eng siRNA and the anti-sEng antibody did not alter the Eng siRNA-driven proliferative effect. Taken together, these results indicate that sEng plays a minimal role in regulating CM proliferation, and suggest that the Eng-mediated inhibition of CM proliferation is primarily accomplished by CM-autonomous signaling through the membrane-bound form of the protein.

**Figure 4.**
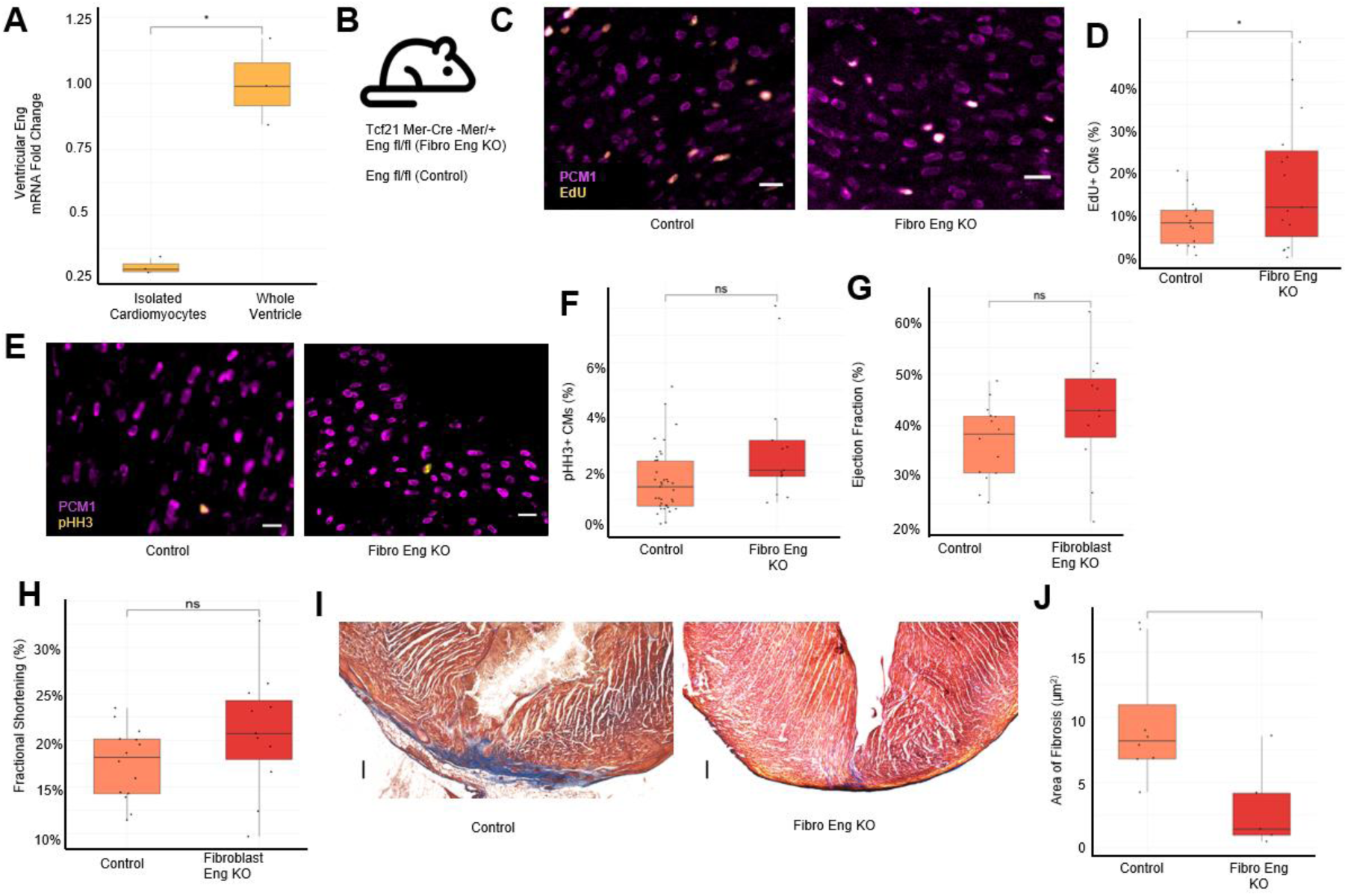
Fibroblast Eng Does Not Inhibit CM proliferation. (A) Relative Eng mRNA expression in isolated CMs vs whole ventricle from 7-day old mice (B) Experimental groups for conditional fibroblast Eng-KO experiments (C) Representative images from control and Fibro Eng KO showing EdU and Pcm1 staining (D) Quantification of control and Fibro Eng KO mice 13 days after injury, N=15/15 (E) Representative images from control and Fibro Eng KO showing pHH3 and Pcm1 staining (F) Quantification of control and Fibro Eng KO mice 7 days after injury, N=37/13 (G) Representative Masson’s Trichrome Staining from control and Fibro Eng KO mice 39 days after injury (H) Quantification of fibrosis of control and Fibro Eng KO mice 39 days after injury, N=8/5 (I) Ejection Fraction and (J) Fractional Shortening of control and Fibro Eng KO mice 38 days after injury, N= 14/11 (K) Quantification of cleavage-furrow localized Aurora B Kinase and Troponin T staining of human CMs after treatment with *Eng* siRNA and/or sEng blocking antibodies (N=288/384 wells).

### Endoglin Modulates Cardiomyocyte Proliferation Through Smad2 Signaling

We next wished to gain a causative understanding of Eng’s effect on CM proliferation. Eng has been shown to signal independently of TGF-β signaling through binding with Zyxin (Zyn) (Conley et al., 2004). However, Zyn siRNA did not reduce the stimulatory effect of Eng siRNA on CM proliferation (Supplemental Figure 5A-B). In addition to Zyn signaling, Eng is known as a TGF-β and BMP signaling co-receptor. To determine a possible effect on TGF-β or BMP signaling, we performed a series of experiments where we knocked-down components of the TGF-β/BMP signaling pathway in conjunction with Eng siRNA and measured their ability to induce S-phase progression of NRVMs using EdU incorporation as the main readout of modulation of proliferation.

Eng has been implicated in the proliferative effects of the Bmp signaling pathway (Jonker, 2014; Saito et al., 2017; Scherner et al., 2007). Eng can bind with Bmp9 and regulate Bmp signaling (Nolan-Stevaux et al., 2012; Núñez-Gómez et al., 2017). Knock-down of Bmp9 resulted in a reduction in CM proliferation with or without Eng siRNA treatment (Figure 5A and Supplemental Figure 5B). An important receptor that mediates Bmp signaling is the type 2 Bmp receptor, Bmpr2, which has been shown to bind Eng (Hiepen et al., 2019; Nolan-Stevaux et al., 2012). Interestingly, depletion of *Bmpr2* alone did not alter CM proliferation, but *Bmpr2* knock-down in conjunction with *Eng* knock-down resulted in higher CM proliferation than *Eng* knockdown alone (p<0.0001), suggesting that signaling through Bmpr2-Eng mediates inhibition of CM proliferation. Smad1 and Smad5 are transcription factors activated downstream of Bmp binding to the Bmp receptor. Knock-down of either *Smad1* or *Smad5* reduced CM proliferation. Surprisingly, simultaneous knockdown of *Eng* with either *Smad1* or *Smad5* robustly increased CM proliferation (Figure 5A and Supplemental Figure 5B). Taken together these data indicate that Bmp signaling is likely involved in regulating CM proliferation, although the involvement of Eng in this pathway is not entirely clear due to opposing effects.

**Figure 5.**
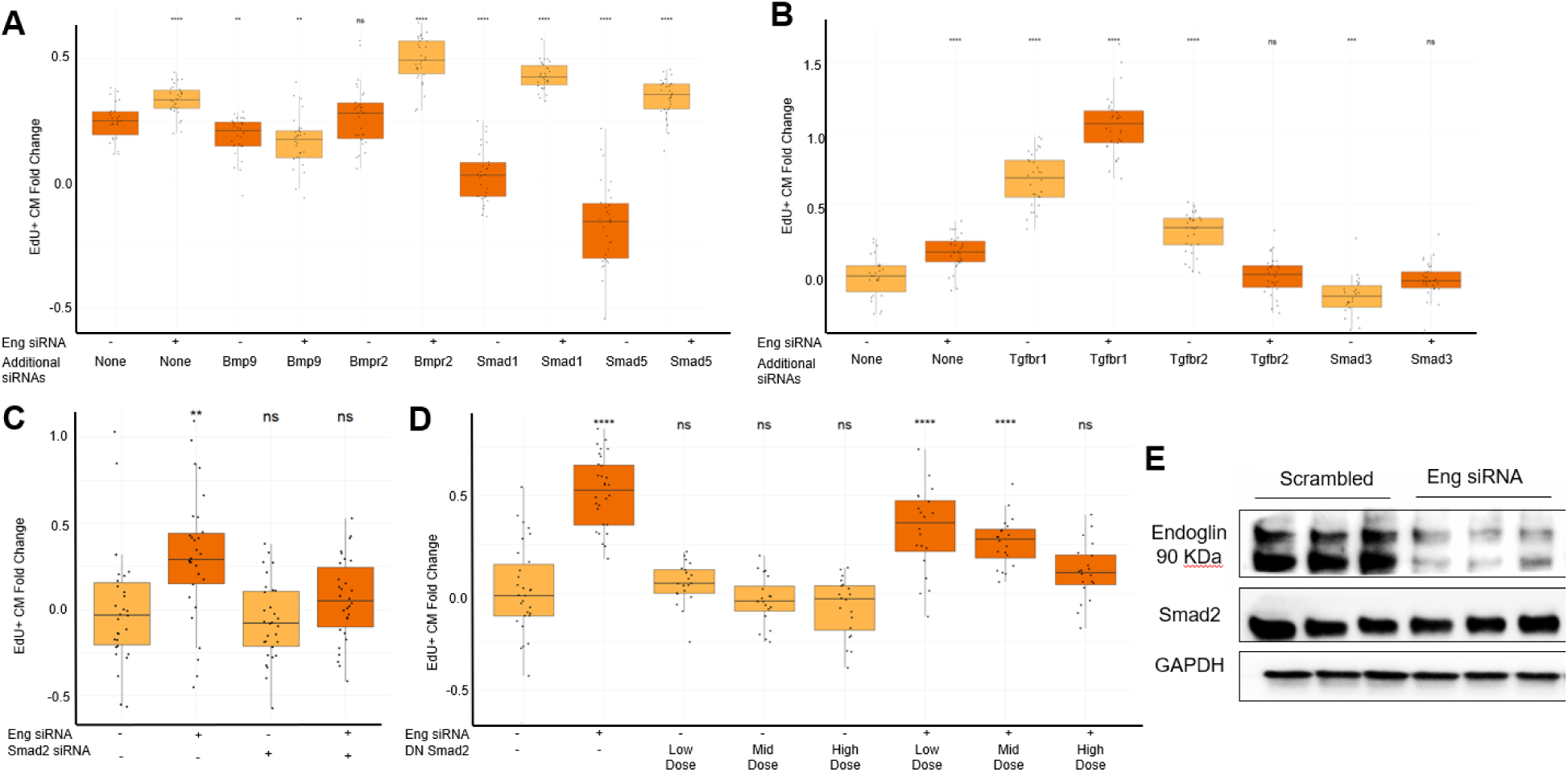
Eng Inhibits CM proliferation in a Smad2 dependent manner. (A) Quantification of EdU+ CMs of NRVMs treated with siRNAs against Eng and/or indicated Bmp signaling pathway members (B) Quantification of EdU+ CMs of NRVMs treated with siRNAs against Eng and/or Tgf-β signaling pathway members (C) Quantification of EdU+ CMs of NRVMs treated with siRNAs against Eng and/or a dominant negative Smad2 adenovirus (D) Western blotting of NRVM protein lysates for Eng, Smad2 and Gapdh treated with scrambled or Eng siRNA (E) Quantification of the Smad2 expression from NRVMs treated with scrambled or Eng siRNA.

The other major branch of the TGF-β signaling pathway that can be influenced by Eng is the canonical TGF-β pathway, where mEng can act as a coreceptor to TGF-β receptors 1 (Tgfbr1) and 2 (Tgfbr2), which leads to activation of Smad2/3 transcription factors. Therefore, we investigated whether Eng functionally interacts with the TGF-β-Smad2/3 pathway in CM proliferation. Depletion of *Tgfbr1* alone enhanced CM proliferation, while combinatorial knockdown of *Tgfbr1* and *Eng* further stimulated CM proliferation (p<0.0001), suggesting potential additive or synergistic effects between Eng and Tgfbr1 (Figure 5B and Supplemental Figure 5D). Interestingly, depletion of *Tgfbr2* alone promoted proliferation, while simultaneous knockdown of *Eng* and *Tgfbr2* resulted in a reduction of CM proliferation to baseline levels (Figure 5B and Supplemental Figure 5D). These results clearly implicate TGF-β receptor signaling in the Eng-mediated negative regulation of CM proliferation. TGF-β signaling can act through both canonical (Smad2/3 dependent) and non-canonical (Smad2/3 independent) signaling pathways. In the non-canonical signaling branch, p38 protein kinase has been shown to inhibit CM proliferation (Engel et al., 2006; Engel et al., 2005). To assess a potential impact of p38 signaling on the Eng-mediated proliferative effects, we used adenoviral overexpression of p38. Moderate and low overexpression p38 alone promoted CM proliferation, albeit at lower levels compared with *Eng* siRNA. However, when combined with *Eng* siRNA, the ability of *Eng* siRNA to enhance proliferation was neutralized with low and moderate p38 overexpression (Supplement Figure 5C). These results suggest that p38 signaling may contribute to the Eng-mediated inhibitory effect on CM proliferation. Finally, within the canonical TGF-β signaling pathway, Smad2 and Smad3 mediate intracellular signaling. *Smad3* depletion alone resulted in a decrease in proliferation while *Smad3* combined with *Eng* siRNA showed similar proliferation rates to scrambled siRNA control conditions (p=0.3692) (Figure 5B and Supplemental Figure 5B). To probe the involvement of Smad2 signaling in the Eng-mediated effect on CM proliferation, we used a dominant negative Smad2 (DN-Smad2) adenovirus. Inhibiting Smad2 signaling did not alter the rate of CM proliferation in conjunction with scrambled siRNA (Figure 5C). However, combination of the DN-Smad2 virus with *Eng* siRNA reduced proliferation rates to control levels in a dose dependent manner. This data indicates that Smad2 signaling is required for the proliferative effect of *Eng* knock-down. To further investigate whether Eng-inhibition of CM proliferation involves direct modulation of Smad2 by Eng, we examined Smad2 expression levels in the setting of *Eng* knock-down. Depletion of Eng did not result in a significant change in Smad2 protein levels (Figure 5D-E). This suggests that Smad2 mediated Eng signaling is likely regulated by Smad2 activation instead of it’s protein levels. Overall this data suggests that the proliferative effect of Eng knockdown requires SMAD2 signaling. Overall, both TGF-β and BMP signaling are likely involved downstream of Eng but the exact mechanism by which this occurs is unclear. Our results implicate Smad2 as a required mediator of the proliferative effects of Eng knockdown.

### Endoglin Inhibits Human CM Proliferation

Finally, we wished to see if *Eng* also inhibits CM proliferation in humans. We performed *Eng* knockdown on human induced pluripotent stem cell (iPSC)-derived CMs. Similar to mouse and rat CMs, human CMs showed enhanced DNA synthesis levels, measured by increased EdU incorporation (Figure 6A-B). Furthermore, we detected increased numbers of human CMs with Aurora B Kinase staining at cleavage furrows indicative of dividing cells in response to *ENG* knockdown (Figure 6C-D). We next sought to identify the mechanism via which Eng mediates CM proliferation. To this end we performed bulk RNA sequencing on human CMs (Supplemental Table 2). Eng siRNA CMs exhibited a significant up regulation in several mitosis related GO terms and a down regulation of multiple GO terms related to muscle contraction (Figure 6E). Kinase enrichment analysis showed an overrepresentation of expression targets for three major cell cycle progression promoting kinases (Cdk1, Pkl1 and AurkB) (Supplemental Table 2). Interestingly, treating human CMs with Eng siRNA did not alter Smad2 transcript levels, which suggests that this interaction is not regulated at the transcript level. Smad2 siRNA treatment also did not alter Eng mRNA levels Supplemental Table 2).

**Figure 6.**
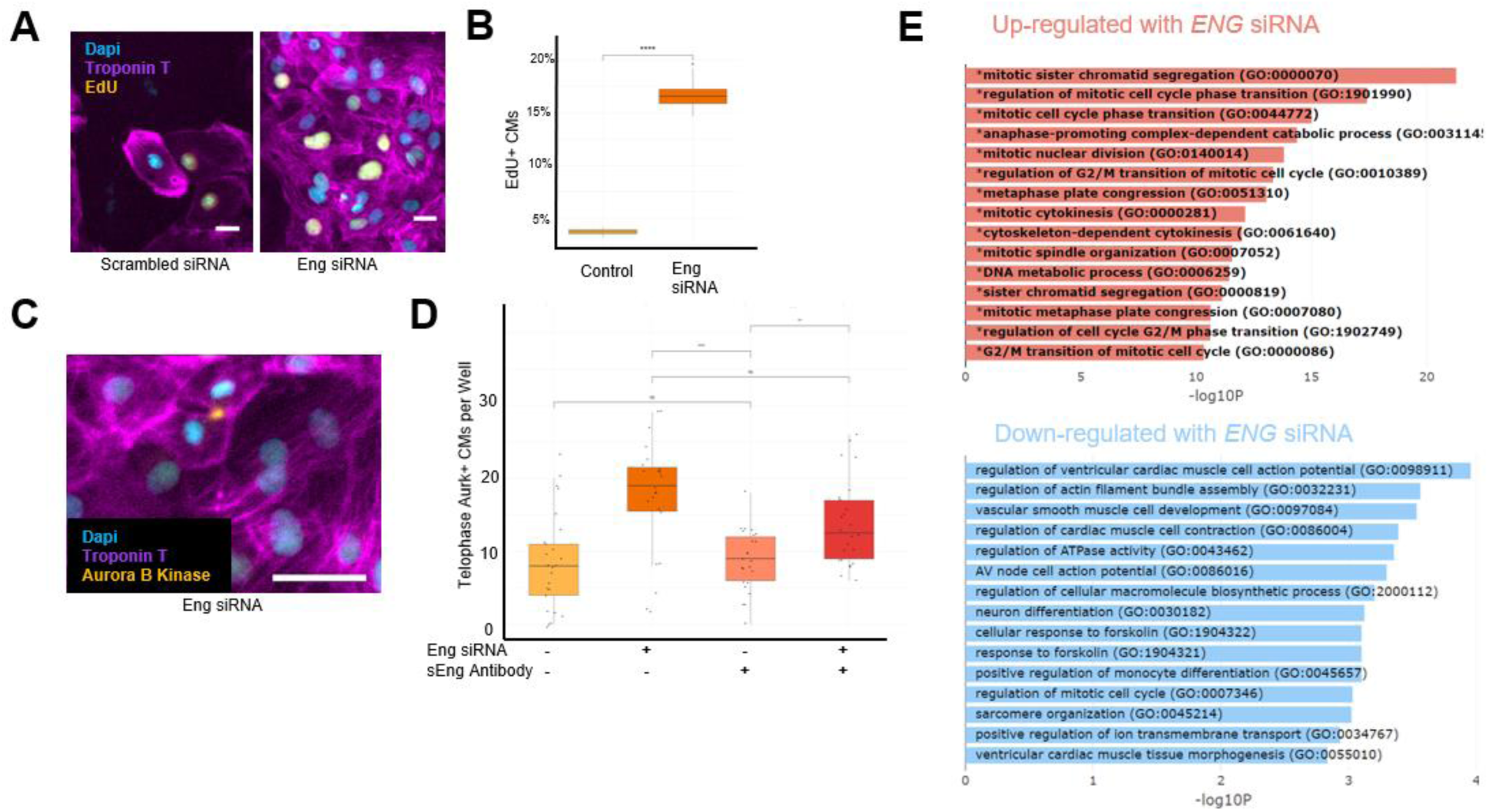
Eng Inhibits Human CM-Proliferation. (A) Representative images of scrambled or Eng siRNA treatment showing EdU incorporation in human CMs. (B) Quantification of EdU positive Human CMs after treatment with scrambled or Eng siRNA. (C) Aurora B Kinase and Troponin T staining in human CMs. (D) Quantification of cytokinetic events (telophase) based on Aurora B Kinase staining between human CMs after treatment with scrambled or Eng siRNA. (E) Significantly enriched GO Terms of human CMs treated with Eng siRNA relative to scrambled siRNA.

## Discussion

Adult CMs are unable to proliferate and divide under both physiological and pathological conditions, such as those after injury. Although control of cell cycle progression has been studied extensively in the setting of cancer development, the heart appears refractory to tumor formation, suggesting that CMs might have additional layers of regulation to control cell cycle entry. In recent years a number of chemicals and genes that can regulate CM proliferation have been identified through unbiased screens or hypothesis-based experimentation. Previous screens have mostly focused on microRNAs or chemical compounds (Diez-Cuñado et al., 2018; Dogan et al., 2018; Magadum et al., 2017; Mills et al., 2019; Uosaki et al., 2013; Woo et al., 2019). These screens identified various compounds that stimulate CM proliferation in cell culture. Beyond these screens, genetic regulators of CM proliferation included both stimulatory ones, such as soluble growth factors, receptors, cyclins, cyclin dependent kinases and transcription factors, as well as inhibitory ones, such as P38 Mapk, Hippo-Salvador, cyclin dependent kinase inhibitors and retinoblastoma proteins (Foglia and Poss, 2016; Mohamed et al., 2018; Tamamori-Adachi et al., 2003). Here, we performed a genome-wide screen to identify additional inhibitors of CM proliferation. Our screen identified known inhibitors of cell cycle progression, such as Cyclin dependent kinase 1a (Cdkn1a) and Retinoblastoma Transcriptional Corepressor Like 1 (Rbl1). In addition to these known inhibitors, our screen identified mainly genes that were not known to play a role in CM proliferation. Knockdown of many of these genes showed enhanced cell cycle activity in cell culture. Interestingly, when we tried to confirm regulation of CM proliferation in vivo, some of the genes identified by our screen appeared to be dispensable for regulation of CM proliferation in vivo. This discrepancy is not unique, as others have observed similar discrepancies between cell culture and in vivo regulation of proliferation (Cutie et al., 2020).

Our screen identified Eng as a novel regulator of CM proliferation. Knock-down of Eng was sufficient to promote both S-phase and M-phase progression in vitro. Furthermore, our results showed enhanced mRNA expression of this proliferative inhibitor in adult CMs, which correlates with the developmental transition from the immature regenerative stage to the mature non-regenerative stage in adulthood. This raised the possibility that Eng might help regulate to this developmental transition. To test this, we deleted Eng from CMs or fibroblasts and subjected neonatal hearts to apical resection. Interestingly, only CM-specific deletion of Eng resulted in enhanced CM proliferation and enhanced cardiac regeneration. However, the Eng mediated proliferative and regenerative phenotype is not as strong as that observed in P1 mice suggesting that Eng is not solely responsible for the developmental transition.

Eng can be expressed as its full-length version where it acts as a membrane-bound TGF-β family coreceptor (mEng), or the extracellular domain can be cleaved to form a soluble peptide with anti-angiogenic effects (sEng). These two forms of Eng have been shown to have diverse, and at times antithetical effects on cellular proliferation (Lebrin et al., 2004; Lin et al., 2013; Núñez-Gómez et al., 2017). Furthermore, sEng can bind circulating growth factors, such as Bmp9 and mediate interaction with their cognate receptor. Eng is best known as a TGF-β-family coreceptor expressed in endothelial cells and important in angiogenesis. The vast majority of cardiac Eng is produced by non-myocardial cells and thus the majority of sEng is produced by other cell types such as fibroblasts. Primary CM cultures typically contain some residual fibroblasts, so it was important for us to distinguish the effects of mEng vs sEng on CM proliferation. CM-specific deletion of Eng resulted in enhanced CM proliferation, improved cardiac function after injury and reduced scar formation. Importantly, when we deleted Eng from cardiac fibroblasts, there was a reduced effect on CM proliferation with no improvement in cardiac function, despite a reduced scar size 1 month after the myocardial injury, indicating that CM mEng is responsible for mediating inhibition of CM proliferation. These results were corroborated by antibody blocking and siRNA experiments using hiPSC-derived CMs, where siRNA against Eng enhanced CM cell division, which was not blocked by an sEng blocking antibody. The human CMs were substantially more proliferative after Eng siRNA both in terms of S-phase progression and cytokinesis. This increase in proliferation is either a result of differences between human and rodent Eng or a result of the fact that the human CM culture might contain lower amounts of sEng due to lower numbers of non-CMs in that culture. There are several Eng targeting therapeutics in clinical trials such as anti Eng monoclonal antibodies (Nolan-Stevaux et al., 2012). Unfortunately, these specifically suppress the sEng not mEng and thus are not ideal to regenerate the myocardium. Our work suggests that CM mEng inhibition could be a promising therapeutic option to repopulate the myocardium.

It is interesting to note is that both CM and fibroblast Eng KO resulted in reduced fibrosis after myocardial injury. This suggests that Eng is important in both myocardial repopulation and either the deposition or reabsorption of fibrosis. Interestingly, the fibroblast Eng KO was equally successful at reducing fibrosis after injury as the CM KO. Furthermore, these results indicate that fibroblasts play an important role in the reduction of cardiac fibrosis. They also support previous work which showed that mice with an Eng haploinsufficiency display less cardiac fibrosis after injury and that soluble Eng expression drives Tgfβ1 mediated collagen deposition (Kapur et al., 2012). Taken together this suggests that cardiac regeneration is promoted by the coordination of repopulation of the myocardium and the resolution of fibrotic scarring.

The ability of Eng to modulate proliferation through both the TGF-β and Bmp signaling pathways is well documented, although the direction of the effect depends on the cell type and context, suggesting the involvement of unknown co-regulators. In mouse endothelial cells Eng can enhance proliferation induced by low doses of TGF-β. Conversely, it can counteract the anti-proliferative effect of high dose TGF-β (Lebrin et al., 2004). Meanwhile, the proliferation of rat myoblasts is inhibited by TGF-β, which can be overcome by Eng overexpression (Letamendía et al., 1998). Eng has been repeatedly shown to modulate proliferation through both TGF-β and Bmp signaling and a good explanation for the observed anti- or pro-proliferative effects is lacking (Núñez-Gómez et al., 2017). The proliferative signaling through which Eng modulates proliferation appears to be very cell type dependent and so it is important to understand it’s effect in CMs.

We explored a possible role of CM Eng signaling in both the BMP and the TGF-β signaling pathways. The Bmp signaling pathway is a potent regulator of proliferation during cardiac development. sEng has been shown to directly bind Bmp9 and the two can complex to form an active signaling molecule (Lawera et al., 2019). Our results indicated reduced proliferation by Bmp9 inhibition itself, with the combination of Eng and Bmp9 siRNA inhibiting CM proliferation to below baseline levels, suggesting a potential role for Bmp signaling in CM proliferation. The ability of Bmp9 inhibition alone to limit CM proliferation suggest that Bmp9 may have a minor role in modulating proliferation independent of Eng. This is further corroborated by our results with inhibition of the mEng binding partner Bmpr2 and the two major downstream effectors Smad1 and Smad5 (Sorensen and van Berlo, 2020). The knockdown of any of these three genes resulted in a decrease in proliferation, independently of Eng inhibition. Interestingly, when Bmpr2, Smad1 or Smad5 inhibition was combined with Eng inhibition we observed a stimulatory effect on CM proliferation. These results indicate that the Bmp pathway is indeed an important regulator of CM proliferation, but the pro-proliferative effects of Eng inhibition do not solely depend on active signaling through the Bmp pathway.

We observed different results for inhibition of members of the canonical Tgf-β pathway. Inhibition of Tgfbr1 and Tgfbr2 alone resulted in enhanced proliferation, while dual inhibition of Tgfbr1/Eng dual inhibition resulted in the strongest enhancement of proliferation. These results suggest that Eng inhibition works in concert with Tgfbr1-mediated inhibition of CM proliferation, and might indicate that combined Eng/Tgfbr1 inhibition could further potentiate the regenerative response, similar to that observed with Wee1 and Tgfbr1 inhibition in conjunction with Cdk4 and Ccnd overexpression in adult mice (Mohamed et al., 2018). The inhibitory effect of Tgfbr2 and Eng on CM proliferation appears to be codependent as indicated by a return to baseline levels after co-treatment. However, the exact nature of this interaction is unclear. Tgbr2 is clearly important for Eng mediated proliferation, but the hierarchy and mechanism of this interaction has yet to be understood. Since Smad2/3 are the major effectors of the canonical TGF-β and activin signaling pathways (Sorensen and van Berlo, 2020), we next assessed whether Eng could signal through these transcription factors. Surprisingly, Smad2 and Smad3 showed divergent effects. Smad3 inhibition by itself resulted in reduced proliferation, while the combined inhibition of Smad3 and Eng restored CM proliferation to baseline levels. A clearer effect on CM proliferation was observed with Smad2 and Eng modulation. Smad2 disruption alone did not alter CM proliferation. However, the application of a dominant-negative Smad2 was able to reduce Eng siRNA CM proliferation levels to baseline levels in a dose dependent manner. This demonstrates that Eng inhibits proliferation in a Smad2 dependent manner. Taken together, these results indicate that Eng inhibits CM proliferation through the suppression of Smad2 signaling or activation. Smad2 can form a variety of transcriptional complexes with distinct transcriptional changes (Lucarelli et al., 2018). It is likely Eng is acting, at least in part, through one of these other complexes. Future work is needed to identify the precise Smad2 signaling complex that modulates proliferation and what the downstream transcriptional targets are. A better understanding of the downstream signaling via which Eng mediates proliferation could also be advantageous for a clinical setting. Global Eng inhibition results in adverse effects on the endothelium and thus unspecific targeting of Eng may also result in adverse phenotypic outcomes(Fernández-L et al., 2006). Instead, a more focused understanding of the downstream signaling of Eng may allow us to identify a therapeutic target for myocyte replacement that is not essential for proper angiogenesis.

In conclusion, our genome-wide screen identified Eng as an inhibitor of CM proliferation and cardiac regeneration. Eng was sufficient to inhibit progression of CMs through all stages of the cell cycle. CM Eng knock-out, but not fibroblast Eng knockout, was sufficient to induce CM proliferation in-vivo after injury and functional regeneration. The ability of Eng to regulate proliferation is dependent upon Smad2 signaling but other members of the TGF-β and Bmp signaling pathways appear to be relevant for mediating cardiomyocyte proliferation as well. An integrated understanding of the various signaling pathways that govern CM proliferation will be an important topic of study going forward. It will also be important to assess if Eng inhibition can restore the CM proliferative effects in adulthood and enhance cardiac regeneration in response to myocardial infarction. Additional regulators of CM proliferation are likely to be discovered, and future screening could be improved by incorporation of in-vivo CRISPR/Cas9 mediated gene deletion after neonatal myocardial injury (VanDusen et al., 2021).

## Supporting information

Key Resources

## Acknowledgments

We would like to thank Chen Chen, Allison Leopold and Delaney Peterson for their help with the histology and Ingrid Bender for her help with adult CM isolation on this manuscript. This work was supported by the resources and staff at the University of Minnesota University Imaging Centers: SCR_020997. D.W.S. was supported by NIH (T32GM113846), J.H.v.B. was supported by grants from the NIH, Regenerative Medicine Minnesota: (RMM 102516 009), Saving tiny Hearts Society and The Hartwell Foundation. R.P. is supported by NIH (R01 AR078571 and R56 AR05529) (R.C.R.P.). B.I.G. was supported by NHLBI (F30 HL151138) and NIGMS (T32 GM008244).

## Author contributions

Conceptualization, D.W.S., J.D.M., J.H.v.B.; Methodology, D.W.S., P.M.T., J.S., J.J.S., J.H.v.B.;Software, P.M.T., J.S., J.J.S.; Investigation, D.W.S., P.M.T., B.I.G., D.Y., M.J.Z., C.G., H.E., K.J., J.H.v.B., Writing, D.W.S., J.H.v.B.; Resources, J.D.M., R.C.R.P.; Funding acquisition, J.D.M., J.J.S., J.H.v.B.; Supervision, T.D.O.C., R.C.R.P., Y.K., J.J.S., J.H.v.B.

## Methods

### Experimental Models and Subject Details

#### Animals

All animal experiments were performed in accordance with the NIH guidelines and approved by the Institutional Animal Care and Use Committee at the Cincinnati Children’s Hospital Medical Center and the University of Minnesota. Fetal murine cardiomyocytes were obtained from time-mated pregnant CD1 mice, *Mus musculus* (Charles River Laboratories). Neonatal rat ventricular cardiomyocytes were obtained from newborn Sprague-Dawley rat pups, *Rattus norvegicus* (Charles River Laboratories). Cdkn1a knockout mice (016565) were obtained from the Jackson laboratory. Endoglin loxP mice, originally generated by the Arthur laboratory (Allinson et al., 2007), were cross-bred with the Myh6-MerCreMer (Myh6-MCM) mouse line obtained from the Jackson laboratory and with the Tcf21-MerCreMer (Tcf21-MCM) mouse line obtained from the Tallquist laboratory (Acharya et al., 2011). Both of these Cre lines have been previously shown be highly efficient at inducing recombination and have negligible leakiness, with comparable tamoxifen doses (Acharya et al., 2011; Yan et al., 2015). The Mapre3 loxP mouse line was obtained from the EMMA mouse repository (EM 07740). The neomycin cassette was deleted by cross-breeding to a germline Flp expressing mouse strain (012930) and the resulting Mapre3^fl/fl^ mouse line was cross-bred with the Myh6-MCM mouse line. Genotyping was done on tail DNA using previously published primer sets for the respective mouse lines. (Z)-4-OH Tamoxifen (60mg/kg) was administered subcutaneously to neonatal pups at postnatal day 1. For experiments including the Myh6-MCM mouse line we used Eng^fl/fl^ X Myh6-MCM and Mapre3^fl/fl^ X Myh6-MCM with tamoxifen as experimental groups and Myh6-MCM with tamoxifen as control. For experiments including the Tcf21-MCM mouse line we used Eng^fl/fl^ X Tcf21-MCM with tamoxifen as the experimental group and Eng^fl/fl^ with tamoxifen as controls. Apical resection was performed at postnatal day 7 as previously described (Mahmoud et al., 2014). We extracted DNA from cardiac tissue of experimental and control animals and performed PCR genotyping to detect genetic recombination of the floxed allele. We only included Cre positive experimental samples that showed recombination. For P21 KO and control animals, hearts were fixed by perfusion fixation with 4% PFA, followed by excision of the heart and digestion with a Collagenase II solution (60mg/ml) for 2h at 37°C as previously described (Yücel et al., 2020). After digestion, the suspension was filtered through a 200µm mesh and CMs were enriched by centrifugation at 100g for 1min. Isolated CMs were further processed for immunostaining.

#### Embryonic Murine Cardiomyocyte Cell Culture

Time-mated pregnant CD1 mice at E17.5 were euthanized by cervical dislocation under isoflurane anesthesia. Embryos were extracted, decapitated and the thorax was opened via sternotomy to expose the heart. The heart was excised and the atria were removed. The murine fetal ventricles were incubated in Trypsin in Hanks Buffered Saline Solution (HBSS, Mg^2+^ and Ca^2+^ free) for 2h at 4°C. The ventricles in trypsin were put at RT for 15min, and Soybean Trypsin Inhibitor (Worthington SIC) in HBSS was added, followed by incubation at 37°C for 15 min. Collagenase dissolved in Leibovitz L-15 was added and incubated at 37°C for 60min with gentle mixing. The ventricles were triturated 10-15 times followed by an additional 15 min incubation at 37°C and additional trituration for 10-15 times. The cells were filtered through a 40µm strainer and pre-warmed M199 with 15% Fetal Bovine Serum (FBS) was added. Cells were pelleted by centrifugation at 1000rpm for 5 min. Cells were resuspended in M199 with 15% FBS,and again pelleted by centrifugation, followed by resuspension in M199 with 2% FBS. Cells were incubated in an uncoated 175cm^2^ tissue culture flask for 1h. After 1h the unattached cells were pipetted off, pelleted by centrifugation and resuspended in M199 with 2% FBS. Cells were counted and diluted to 80,000 cells/ml and 50µl cell suspension was added to each well of a 0.1% gelatin-coated 384-well plate. After seeding cells, we added 10µl lentivirus generated from bacterial stocks of the MISSION TRC1 library to each well, except for control wells. We used Shc002 lentivirus as a negative control and Myc overexpressing adenovirus as positive control. After addition of viral particles, the plate was quickly spun and allowed to settle at RT for 60min, after which the plate was incubated at 37°C overnight. The next morning, 50µl media was aspirated and 50µl fresh M199 with 2% FBS was added. Two days later the media was exchanged again. The next morning, 50µl media was aspirated and 50µl M199 with 2% FBS and 10µM 5-ethynyl-2’-deoxyuridine was added. Twenty-four hours later the media was aspirated and cells were incubated in 4% paraformaldehyde for 10min, followed by 2 washes in Phosphate Buffered Saline (PBS). For the confirmation experiments, we generated new lentivirus for all clones that were considered hits in the primary screen. The procedure for the confirmation experiments followed the same procedure as above, but instead of using 384 wells cardiomyocytes were now plated in 96 wells plates, we seeded 10,000 cells per well, and we used 40µl lentivirus per well.

#### Rat Cardiomyocyte Cell Culture

Rat CMs were isolated one day after birth as previously described using Trypsin and Collagenase digestion (Toraason et al., 1989). After preplating to enrich for CMs, we seeded 35,000 CMs per well on a 96 well plate in 10% FBS. Rat CMs were transfected with siRNAs (see Major Resources document) and lipofectamine RNAi Max for 24 hours in Opti-MEM, with no serum. Alternatively, cells were treated with 1.5, 3 or 6ul of either 2.20 × 10^11^ pfu/ml B-Gal, dominant negative Smad2 or p38 MAPK Adenoviruses for 12 hours, with no serum. Cells were treated with 10µM EdU between 72 and 96 hours after plating, followed by fixation with 4% paraformaldehyde. All NVRM experimental conditions were replicated from independent isolations and had an N=30 wells with the following exceptions: Supplemental Figure 5C (N=20), Figure 5C (N=20).

#### Maintenance of Human Induced Pluripotent Stem Cells (hiPSCs)

Cells were maintained in mTeSR™-1 medium (STEMCELL Technologies) on growth factor-reduced Matrigel (∼8μg/cm^2^) at 5% CO_2_ and 37°C under normoxia conditions. Cells were singularized with Accutase for 3-5 minutes and passaged every 4 days at a density of 1.25 × 10^4^/cm^2^ with the addition of 10μM ROCK inhibitor for the first 24 hours after each passage. Cell lines were routinely tested for mycoplasma contamination every 3 months.

#### Cardiac Differentiation by the Wnt Modulation Method

After passaging freshly thawed hiPSCs three times in single cell suspension using standard culture methods, CMs were generated using the Wnt modulation protocol as previously described (Lian et al., 2012). This protocol is known to produce predominantly ventricular-like CMs. Briefly, hiPSCs were cultured in 12-well plates to a confluency of 70-85% over 3 days in growth factor-reduced Matrigel (∼22μg/cm^2^) upon which they were then induced with 6.5–8.5μM CHIR 99021 in RPMI1640 supplemented with B27 (minus insulin). Exactly 48 hours later, the media was completely removed and replaced with 7.5μM IWP-2 in RPMI1640 supplemented with B27 (minus insulin) while ensuring that the plates did not remain outside the incubator for more than 5 minutes at a time. Following another 48-hour period, the media is removed and replaced with RPMI1640 supplemented with B27 (minus insulin) on day 4 of the protocol. Beginning on day 6, the media was changed to RPMI1640 supplemented with B27 (plus insulin) and changed every 48 hours. Beating was observed between day 8–12 of the differentiation protocol.

#### Enrichment of Human Cardiomyocytes

HiPSC-derived CMs were purified using lactate to induce a metabolic switch, as previously described (Burridge et al., 2014; Tohyama et al., 2013). Briefly, on day 15 of the differentiation protocol, hiPSC-CMs were incubated with 0.25% Trypsin in EDTA for 20 minutes, singularized using a P1000 pipette, and quenched with three times the volume of trypsin in filtered RPMI1640 with 20% FBS. Cells were centrifuged at 300*g* for 5 minutes and resuspended in RPMI1640 supplemented with B27 (plus insulin) and 10μM ROCK inhibitor. hiPSC-CMs were then seeded onto two 6-well plates per every 12-well plate (∼2.6x dilution by surface area). Forty-eight hours later, the media was changed to lactate media: RPMI1640 without D-glucose, 5mM sodium DL-lactate, 213μg/mL L-ascorbic acid 2-phosphate, and 500μg/mL O. Sativa-derived recombinant human albumin Media was changed every 48 hours for a total of 2 to 6 days upon which the media was reverted to RPMI1640 with B27 (plus insulin) supplement. On day 21-22 of the protocol, hiPSC-CMs were singularized following a similar method as described above and were seeded onto black with clear flat bottom 96-well plates at a density of 6.25 × 10^5^/cm^2^ in growth factor-reduced Matrigel (∼22μg/cm^2^). cTnT staining and flow cytometry was used to confirm that all CMs were of 95% or greater purity (Tohyama et al., 2013).

Human CMs were transfected with siRNAs and lipofectamine RNAi Max for 24 hours in Opti-MEM following the manufacturer’s instructions. In addition to siRNAs, we treated human CMs with anti s-Eng antibodies (TRC105) for an additional 48 hours after transfection. Cells were treated with 10µM EdU between 72 and 96 hours after initial plating, after which cells were fixed with 4% paraformaldehyde and processed for immunocytochemistry. RNA for Human CMs sequencing was also isolated using TRIzol. Human CM sequencing experiments had an N=4 for all conditions and all other experiments had an N=30 per experimental condition.

#### Immunofluorescence, Imaging, and cell quantification

For the genome-wide screen each well was stained for incorporated EdU, troponin T and DAPI. EdU Staining was performed as previously described (Yücel et al., 2020). Briefly, fixed cells were permeabilized with 0.5% Triton X-100 in PBS for 20min, followed by blocking with 3% Bovine Serum Albumin in PBS twice for 10min. EdU staining was performed using click chemistry with the following reaction parameters: 20mM Tris.HCl pH8.5, 4mM CuSO_4_, 10µM Sulforhodamine B Azide, 20mg/ml L-Ascorbic acid at RT for 20min. Each well was washed twice in PBS with 0.1% Triton X-100, followed by incubation with primary antibodies against Troponin T (13-11, Thermo Scientific, both screens) and phospho Histone H3 (sc-8656, confirmation only) in PBS with 0.1% Triton X-100 and 0.1% BSA for 1h at RT. Following incubation with primary antibodies, each well was washed twice in PBS with 0.1% Triton X-100 followed by incubation with secondary antibodies and DAPI. Images were acquired on a Molecular Devices ImageXpress for the primary screen and a Biotek Cytation 3 for the confirmation screen. For ImageXpress imaging, we imaged 18 sites per well of a 384 well plate and for Cytation3 imaging, we imaged 56 sites per well of a 96 well plate. We followed a similar procedure for immunohistochemistry on tissue sections, with the addition of a blocking step (10% donkey serum) prior to primary antibody incubation. A complete list of primary and secondary antibodies can be found in the major resources document. Immunofluorescent mouse tissue sections were imaged on a Axio Observer Z1 (Zeiss), rat, human and fetal mouse CMs were imaged on a Cytation 3 (Biotek) or a Ti-E (Nikon). EdU and pHH3 cardiomyocytes were quantified automatedly using an adapted version of our previously published Cell Profiler pipeline (Bass et al., 2012; Carpenter et al., 2006). Mouse tissue sections that were stained for Aurora B Kinase were imaged using a confocal Nikon A1R microscope. CMs stained for Aurora B Kinase were manually counted and scored by a blinded researcher. Mouse hearts were fixed and stained using Masson’s Trichrome Staining according to manufacturer’s instructions. Trichrome stained sections were imaged using an Axio Imager M2 (Zeiss) and the area of fibrosis was quantified using ImageJ by a blinded researcher. Scale bars are 20μm with the exception of figures (3J,4H) in which they are 100μm and Figure 1 in which they are 50μm.

#### mRNA Quantification

Ventricles of 1- and 7-day old mice were dissected and serially digested in 0.175% Trypsin for a total of 80 minutes. CMs were further purified with a 90-minute preplating on uncoated dishes, followed by collecting the non-adherent cells. Adult mouse CMs were isolated as previously described using retrograde perfusion digestion with Collagenase II (O’Connell et al., 2007). Total RNA was isolated using TRIzol and first strand cDNA was synthesized using 2µg of RNA with SuperScriptIV, both following the manufacturer’s recommendations. qPCR analysis was performed using a Quant Studio 6 Flex using iTaq with RNA polymerase subunit 2 used as a control gene(Radonić et al., 2004). Gene expression was calculated using the delta-delta CT method(Zhang et al., 2013). A list of primers is available in Supplemental Key Resource Table.

#### Western Blot

Rat CMs were transfected with siRNAs (scrambled or Eng) using lipofectamine RNAi Max for 24 hours. Protein extracts were quantified by BCA and normalized, and proteins were separated on a 12% acrylamide gel by SDS-PAGE, followed by blotting on nitrocellulose membranes. Equal protein loading control was verified by Ponceau staining. Membranes were blocked and incubated with primary and secondary antibodies, followed by imaging. Quantification of band intensity was performed using ImageJ relative to GAPDH staining of the corresponding protein sample (Abràmoff et al., 2004).

#### RNA-sequencing

Paired-end libraries of human ventricular tissues using the Illumina platform were retrieved from the gene expression omnibus (GEO) database and the ENCODE consortium (Consortium, 2012; Davis et al., 2018). Raw FASTQ files were obtained for 22 fetal (He et al., 2016; Kuppusamy et al., 2015; Pervolaraki et al., 2018; Spurrell et al., 2019; Szabo et al., 2015; Zhang et al., 2019) (9–28 weeks old), and 34 adult (Asmann et al., 2012; Barbosa-Morais et al., 2012; Churko et al., 2018; Derrien et al., 2012; Johnson et al., 2018; Kuppusamy et al., 2015; Lin et al., 2014; Spurrell et al., 2019; Yang et al., 2014) (30–77 years old) samples. The following GEO Terms were used for Fetal: E-MTAB-7031, GSE106688, GSE35585, GSE62913, GSE64283 and Adults: E-MTAB-7031, GSE106688, GSE35585, GSE62913, GSE64283. Data alignment and comparison of expression was analyzed using the CHURP pipeline (Baller et al., 2019) at the University of Minnesota Genomics Center (UMGC). 2 x 50-150bp FASTQ paired-end reads for 56 samples (28.3 million reads average per sample) were trimmed using Trimmomatic (v0.33) enabled with the optional “-q” option; 3bp sliding-window trimming from 3’ end requiring minimum Q30. Quality control on raw sequence data for each sample was performed with FastQC. Read mapping was performed via HISAT2 (v2.1.0) (Kim et al., 2019) using the human genome (GRCh38.97) as reference. Gene quantification was done via Feature Counts for raw read counts. Differentially expressed genes were identified using the EdgeR (negative binomial, R programming) feature in CLCGWB using raw read counts. We filtered the generated list based on a minimum 2x absolute fold change and FDR corrected *p* < 0.05. Human CM RNA sequencing, including differential gene expression, GO Term analysis and kinase enrichment analysis was performed as described (Torre et al., 2018) siRNA knockdown of Eng was confirmed by sequencing.

#### Echocardiography

To measure left ventricular function after apical resection, we used a Vevo 2100 Fujifilm (VisualSonics, Toronto, CA) ultrasound imaging system to perform echocardiography. The data was collected by a researcher blinded to the genotype of the mice. Mice were anesthetized with 2% isoflurane and affixed in a supine position to a heated platform, while instrumented with a temperature probe and ECG electrodes. Ultrasound measurements were performed with a MS550D linear array ultrasound probe operating between 22-55 MHz. The ultrasound probe was first placed across the mouse thorax to obtain a parasternal long axis view of the heart, and to capture two-dimensional and M-mode cine. The ultrasound probe was then rotated 90 degrees to obtain a parasternal short axis view and to capture two-dimensional and M-mode cine. Echocardiography images were analyzed using the Vevo Lab software by a blinded researcher. Left ventricular ejection fraction was calculated using the modified Quinones method from M-mode measurements of at least 3 different cardiac cycles from either the parasternal long axis or short axis views, while fractional shortening was calculated based on systolic and diastolic dimensions.

#### Statistical Analysis

For the genome-wide screen we calculated a Z’ of 0.5 based on fetal murine CM EdU incorporation rates from negative (shc002) and positive (Myc) controls. Any condition that showed a Z factor of 3 above the negative control (>5% normalized EdU positivity) was considered a potential hit. All images from potential hits were manually inspected to verify accuracy of the automated imaging analysis pipeline. Only hits that were deemed correct were kept for further analysis. For all subsequent experiments, normalization of results was performed to their appropriate control and statistical significance was analyzed using either a Student’s t-test or One or Two-way ANOVA (P < 0.05) with a Tukey’s post-hoc Test. NS indicates not significant, *p<0.05, **p<0.01, ***p<0.001, **** p<0.0001 as compared to the control group or left most bar. All presented data points represent a unique biological replicate, as defined as an individual animal for in-vivo experiments or a distinct well of cells for in-vitro experiments.

