## Supplemental Figures for "Myocardial Endoglin Regulates Cardiomyocyte Proliferation and Cardiac Regeneration"

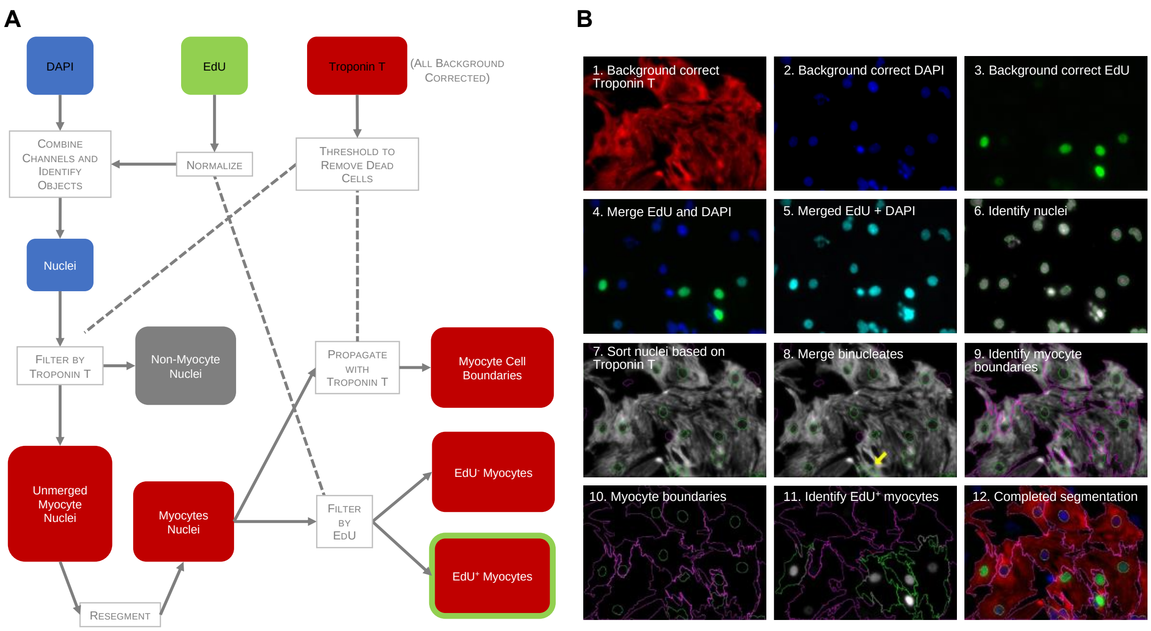


**Supplemental Figure 1. Image Processing Algorithm**
(A) The pipeline used to automatically process images and quantify level of CM proliferation as EdU positive percentage (B) Representative example of sequential steps of the imaging processing algorithm

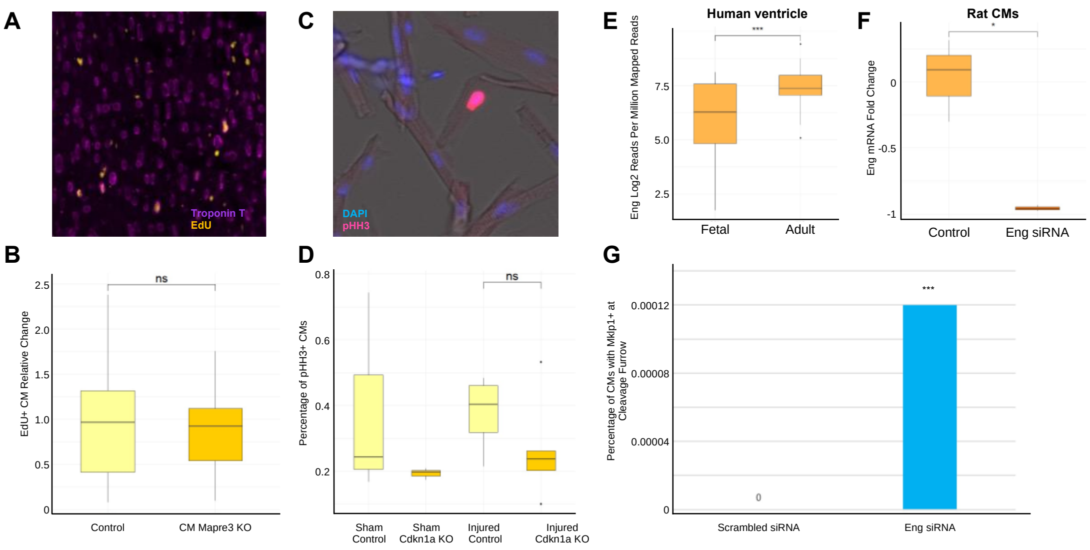


**Supplemental Figure 2. Not all in vitro inhibitors of CM proliferation regulate CM proliferation in vivo**

(A) Representative image of EdU and Pcm1 staining to quantify proliferative cardiomyocytes after apical resection (B) Quantification of EdU positive CMs from control and CM Mapre3 KO mice 13 days after apical resection injury (N=15/18) (C) Representative image of pHH3 staining on isolated CMs (D) Quantification of pHH3 positive CMs from control and Cdkn1a KO mice 7 days after sham or apical resection injury (N=3/3/4/5) (E) Quantification of relative Eng mRNA abundance in human fetal and adult ventricles (F) Quantification of relative Eng mRNA abundance after scrambled or Eng siRNA treatment of rat CMs (G) Quantification of Mklp1 and Troponin T staining of rat CMs after treatment with scrambled or Eng siRNA

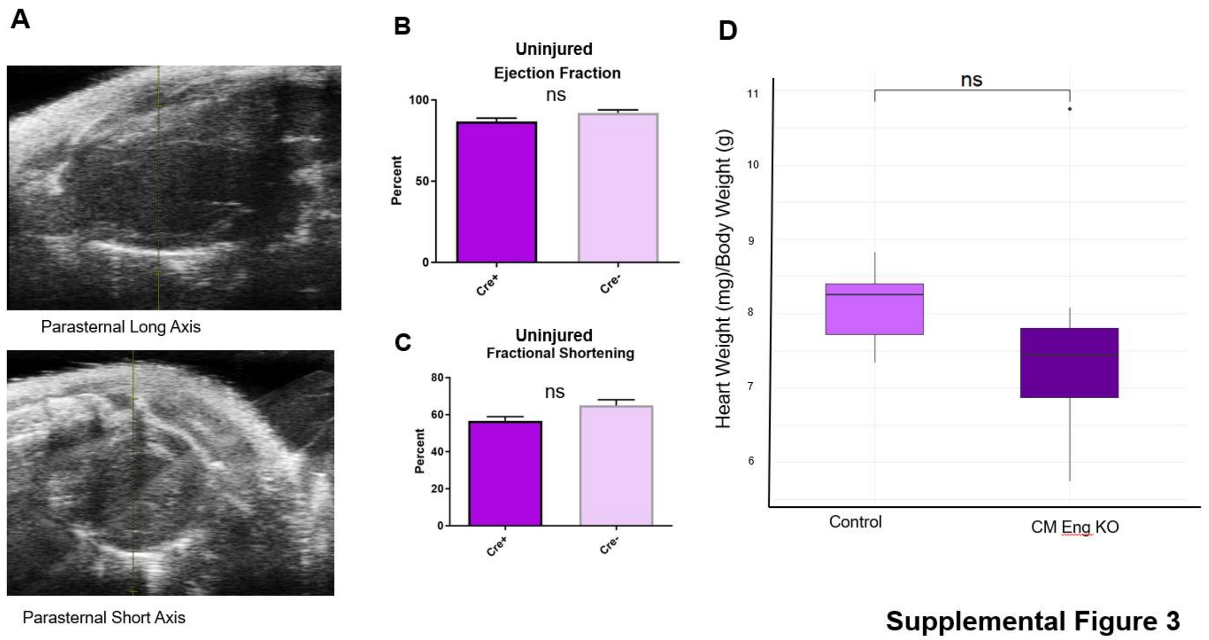


**Supplemental Figure 3**

**Supplemental Figure 3. Representative Echocardiography and CM Eng KO does not significantly alter heart weight to body weight**
(A) Example images of the echocardiography performed on mice 38 days post injury to measure cardiac function (B-C) Quantification of ejection fraction and fractional shortening respectively performed on uninjured Eng fl/fl mice with and without Myh6 Cre. (C) Quantification of the heart weight to body weight ratios of control and CM Eng KO mice 38 days after injury.

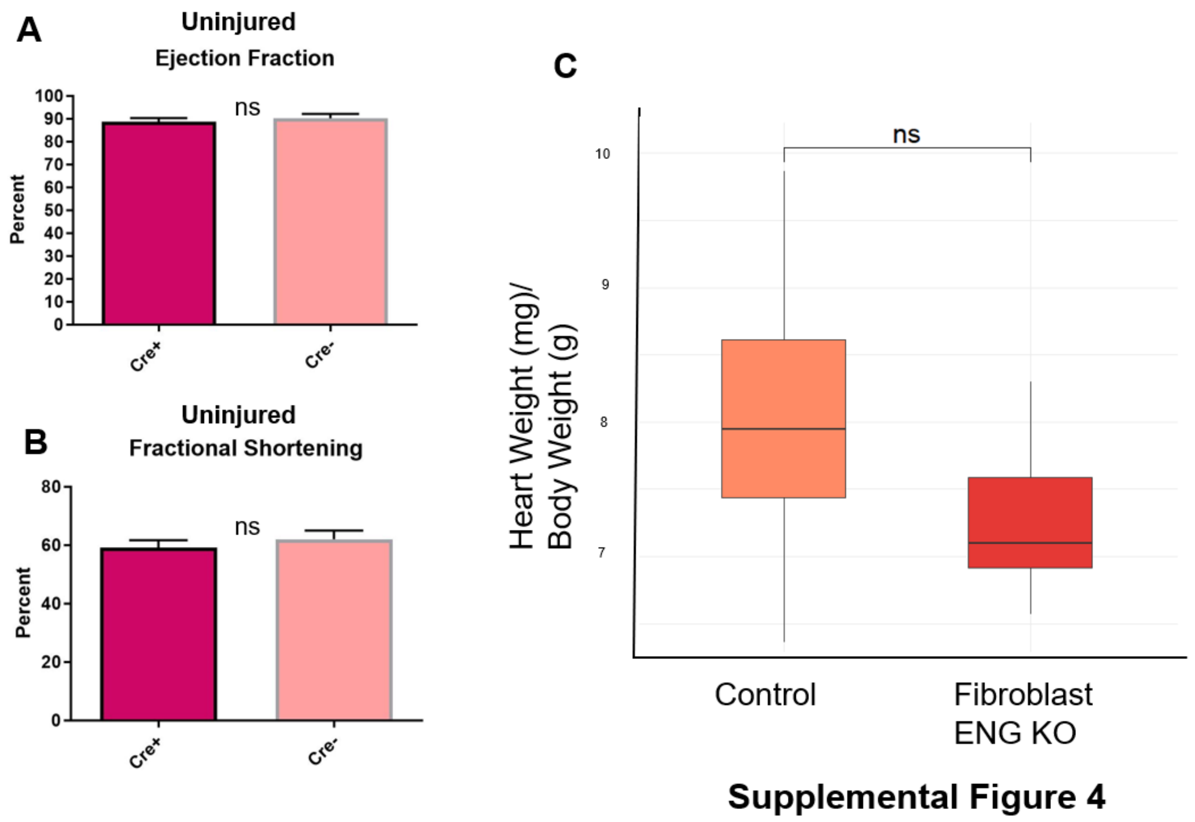


**Supplemental Figure 4. Fibro Eng KO does not significantly alter heart weight to body weight**

(A-B) Quantification of ejection fraction and fractional shortening respectively performed on uninjured Eng fl/fl mice with and without Myh6 Cre. (C)Quantification of the heart weight to body weight ratios of control and Fibro Eng KO mice 38 days after injury.

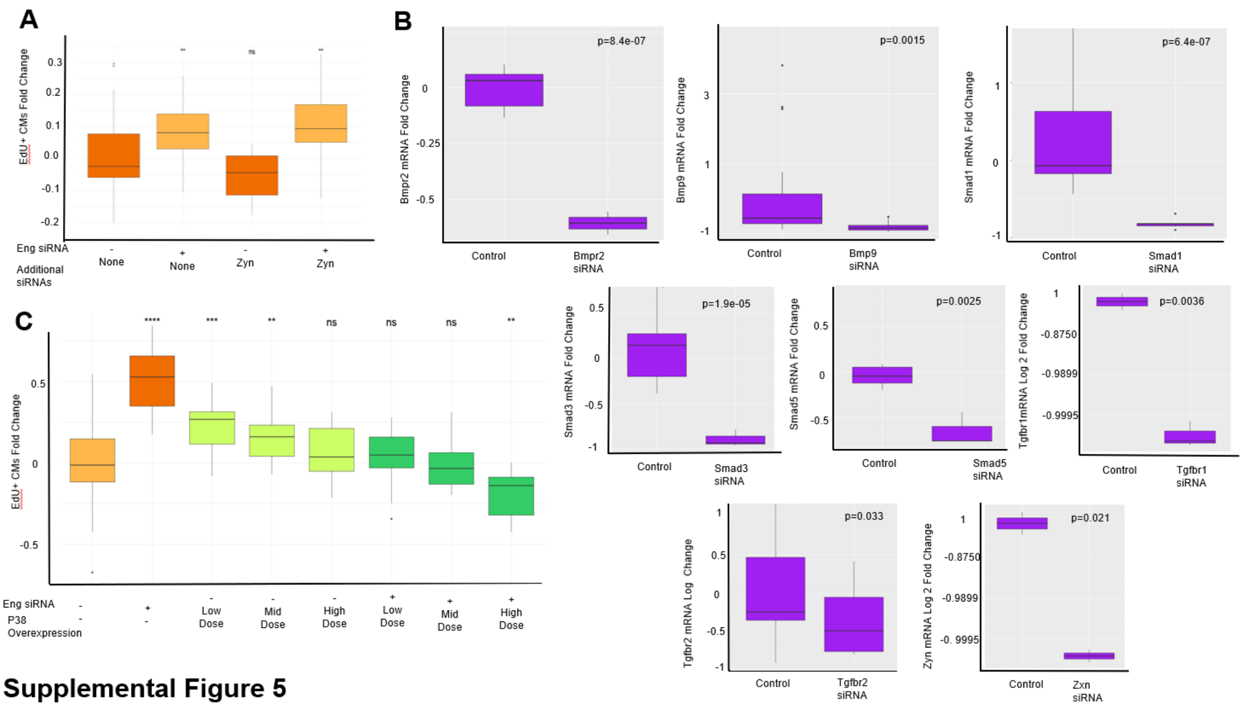


**Supplemental Figure 5**

**Supplemental Figure 5. Verification of mRNA level knockdown after siRNA treatment and effect of Zyn siRNA and p38 on CM proliferation**

(A) Quantification of EdU+ CMs of rat CMs treated with siRNA against Eng and/or Zyn. (B) Relative mRNA expression of rat CMs treated with scrambled Eng related siRNA (C) Quantification of EdU+ CMs of rat CMs treated with siRNAs against Eng and/or p38 expressing adenoviral particles.
