## Supplementary material for "Myocardial Endoglin Regulates Cardiomyocyte Proliferation and Cardiac Regeneration": Key Resources

**Key Resources Table**

| REAGENT or RESOURCE | SOURCE | IDENTIFIER |
| --- | --- | --- |
| **Antibodies** | | |
| Aurora B Kinase Antibody | Abcam | ab2254 |
| Endoglin Antibody TRC105 | Creative Biolabs | AB-232CL |
| Endoglin Antibody MJ7/18 | BD Biosciences | 550546 |
| GAPDH Antibody 6C5 | Santa Cruz Biotechnology | sc-32233 |
| Histone H3 (phospho S10) Antibody | Santa Cruz Biotechnology | sc-8656 |
| Histone H3 (phospho S28) Antibody | Abcam | ab10543 |
| MKLP1 Antibody | Abcam | ab174304 |
| PCM1 Antibody | Sigma-Aldrich | HPA023374 |
| Smad2 Antibody | Cell Signaling | 5339S |
| Troponin T Antibody 19C7 | Novus Biologicals | NB110-2546 |
| Troponin T Antibody 13-11 | Thermo | MS295P |
| **Chemicals, peptides, and recombinant proteins** | | |
| 4',6-Diamidino-2-Phenylindole, Dilactate (DAPI) | Life Technologies | D3571 |
| 5-Ethynyl-2′-deoxyuridine (EdU) | Santa Cruz Biotechnology | sc-284628A |
| 5-Ethynyl-2′-deoxyuridine (EdU) | Carbosynth | NE08701 |
| B27 | Gibco | 17504044 |
| Bovine Serum Albumin | Fisher | bp1605 100 |
| CHIR99021 | Cayman Chemical | 13122 |
| Collagenase, Type 2 | Worthington | LS004176 |
| D-Glucose | Life Technologies | 15023021 |
| DL-lactate | Sigma-Aldrich | L4263 |
| DMEM | Cytiva | SH30243.01 |
| Donkey Serum | Sigma Aldrich | D9663 |
| Fetal Bovine Serum (FBS) | Sigma-Aldrich | 12103C |
| IWP-2 | Cayman Chemical | 13951 |
| L-ascorbic acid 2-phosphate | Sigma-Aldrich | 1713265-25-8 |
| Lipofectamine RNAi Max | Thermo | 13778150 |
| Matrigel | Corning | 354277 |
| Medium 199 | Corning | 10-060-CV |
| PBS | Corning | 21-040-CV |
| O. Sativa-derived recombinant human albumin | Sigma-Aldrich | 70024-90-7 |
| Opti MEM | Thermo | 51985034 |
| Paraformaldehyde | Electron Microscopy Sciences | 15714 |
| ROCK inhibitor | Cayman Chemical | Y-27632 |
| RPMI1640 | Gibco | 11-875-101 |
| TRIzol | Life Technologies | 15596018 |
| Trypsin | Difco | 215240 |
| **Critical commercial assays** | | |
| BCA Protein Assay | Thermo Scientific | PI23225 |
| Cardiomyocyte Isolation Kit | Worthington | LK003303 |
| iTaq | Bio-Rad | 1725124 |
| Trichrome Stain Kit (Modified Masson’s) | ScyTek Laboratories | TRM-1-IFU |
| SuperScriptIV | Thermo | 18090010 |
| **Experimental models: Organisms/strains** | | |
| Mice: C57Bl/6j | Jax #000664 | https://www.jax.org/strain/000664 |
| Mice: CD-1 | Charles River | https://www.criver.com/products-services/find-model/cd-1r-igs-mouse |
| Mice: Eng (fl/fl) | Allinson et al., 2007; | NA |
| Mice: Flpo | Jax #012930 | https://www.jax.org/strain/012930 |
| Mice: Mapre3 (fl/fl) | EMMA ID 07740 and this paper | https://www.infrafrontier.eu/search?keyword=mapre3 |
| Mice: Myh6-MCM | Sohal et al., 2001 | https://www.jax.org/strain/005657 |
| Mice: p21(-/-) | Jax #016565 | https://www.jax.org/strain/016565 |
| Mice: Tcf21 (MCM/+) | Acharya et al., 2011 | NA |
| Rat: Sprague Dawley | Charles River | https://www.criver.com/products-services/find-model/cd-sd-igs-rat |
| **Software and algorithms** | | |
| Cell Profiler | Carpenter et al., 2006 | https://cellprofiler.org/ |
| Cell Profiler Script for In Vitro CM Proliferation Calling | This Paper |  |
| CLCGWB | Qiagen | https://digitalinsights.qiagen.com/products-overview/discovery-insights-portfolio/analysis-and-visualization/qiagen-clc-genomics-workbench/ |
| EdgeR | Robinson, et al., 2010 | https://bioconductor.org/packages/release/bioc/html/edgeR.html |
| ImageJ | Abràmoff et al., 2004 | https://imagej.nih.gov/ij/ |
| HISAT2 | Kim, et al., 2019 | http://daehwankimlab.github.io/hisat2/ |
| Imagej Script for In Vivo CM Proliferation Calling | This Paper |  |
| Trimmomatic | Bolger, et al., 2014 | http://www.usadellab.org/cms/?page=trimmomatic |
| **Oligonucleotides** | | |
| See attached List |  |  |
| **siRNAs** | | |
| Human Eng siRNA | Thermo | 4392420 |
| Rat Bmpr2 siRNA | Thermo | s138926 |
| Rat Eng siRNA | Thermo | s176002 |
| Rat Smad1 siRNA | Thermo | s130428 |
| Rat Smad3 siRNA | Thermo | s130317 |
| Rat Smad5 siRNA | Thermo | s133234 |
| Rat TGFBR1 siRNA | Thermo | s131792 |
| Rat TGFBR2 siRNA | Thermo | s135791 |
| Rat Zyn siRNA | Thermo | s137820 |
| Silencer™ Select Negative Control No. 1 siRNA | Thermo | 4390843 |
